# Crystal Structure of Human Nocturnin Catalytic Domain

**DOI:** 10.1101/330514

**Authors:** Michael A. Estrella, Jin Du, Alexei Korennykh

## Abstract

Nocturnin (NOCT) helps the circadian clock to adjust metabolism according to day and night activity. NOCT is upregulated in early evening and it has been proposed to serve as a deadenylase for metabolic enzyme mRNAs. We present a 2.7-Å crystal structure of the catalytic domain of human NOCT. Our structure shows that NOCT has a close overall similarity to CCR4 deadenylase family members, PDE12 and CNOT6L, and to a DNA repair enzyme TDP2. All the key catalytic residues present in PDE12, CNOT6L and TDP2 are conserved in NOCT and have the same conformations. However, we observe substantial differences in the surface properties of NOCT, an unexpectedly narrow active site pocket, and conserved structural elements in the vicinity of the catalytic center, which are unique to NOCT and absent in the deadenylases PDE12/CNOT6L. Our work thus reveals the structure of an intriguing circadian protein and suggests that NOCT has considerable differences from the related deadenylases, which may point to a unique cellular function of this enzyme.

## Introduction

NOCT is a ∼ 50 kDa eukaryotic phosphodiesterase exhibiting an unusual regulation of expression. Nocturnin levels peak in early evening according to the internal circadian clock^1^. In Drosophila, the NOCT gene is called *curled* (curled wing phenotype)^2^. This phenotype is a known signature of a metabolic rather than a developmental defect^2^. Early studies using recombinant protein from *Xenopus laevis* suggested that NOCT can cleave poly-A RNA^3^, leading to the model that NOCT is a deadenylase destabilizing the mRNAs of metabolic enzymes^4^.

NOCT belongs to a multifunctional protein family. Some NOCT homologues are RNA deadenylases^5,6^. However, NOCT is also similar to the DNA repair enzymes such as APE1^3,7^. The closely related deadenylases are CNOT6L^6^ (a mammalian ortholog of the main yeast deadenylase, CCR4) and PDE12 (a mitochondrial deadenylase required for maturation of mitochondrial tRNAs^5^). In contrast to NOCT, none of the related family members are regulated by circadian clock, suggesting that NOCT has a unique and non-redundant function in regulating metabolism. To begin to understand the structural and molecular basis that underlies this cellular function of NOCT, we determined the crystal structure of the catalytic domain from the human protein.

## Results

### Structure of human NOCT reveals catalytic center and fold similarity to PDE12, CNOT6L, and TDP2

We cloned and purified human NOCT and conducted crystallization of both, full-length protein and truncated variants that contain the catalytic domain. The catalytic domain (residues 122-431) produced crystals that diffracted to ∼ 2.7 Å (Table 1). The resulting NOCT structure revealed a globular protein with α/β sandwich fold shared by multiple phosphodiesterases^6^ (Fig. 1A-B). As has been expected based on the sequence similarity, the NOCT structure is overall similar to the structures of PDE12 and CNOT6L^3,6^ (Fig. 2A). 3D homology search using DALI server^8^ revealed that NOCT has nearly the same degree of similarity also with TDP2, a DNA repair enzyme that resolves topoisomerase stalls by hydrolyzing the phosphodiester link between DNA and tyrosine^9^.

**Table 1.**
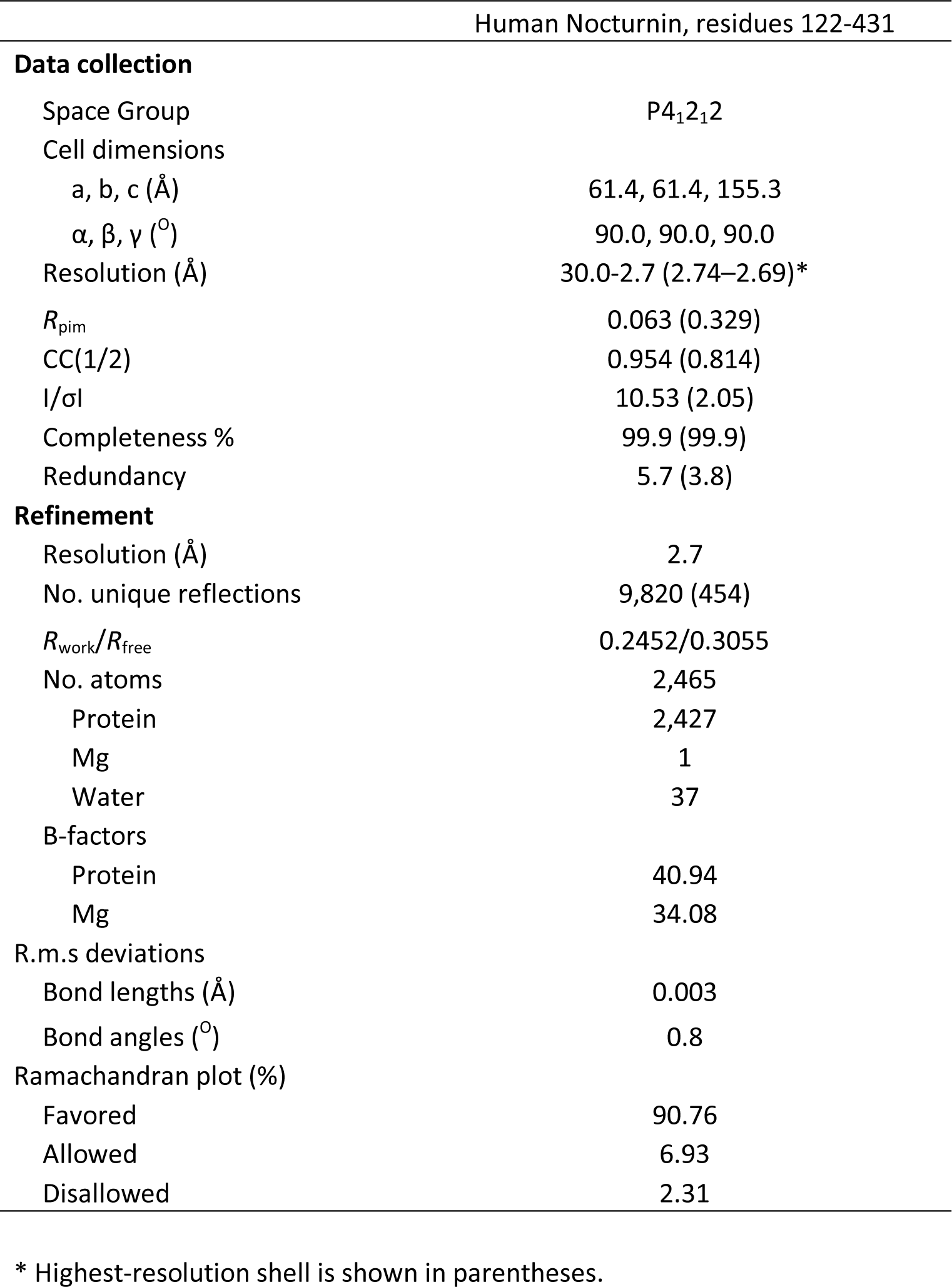
Data collection and refinement statistics.

| Human Nocturnin, residues 122-431 |  |
| --- | --- |
| **Data collection** |  |
| Space Group | P4<sub>1</sub>2<sub>1</sub>2 |
| Cell dimensions |  |
| a, b, c (Å) | 61.4, 61.4, 155.3 |
| $\alpha$, $\beta$, $\gamma$ (°) | 90.0, 90.0, 90.0 |
| Resolution (Å) | 30.0-2.7 (2.74–2.69)* |
| $R_{\text{pim}}$ | 0.063 (0.329) |
| CC(1/2) | 0.954 (0.814) |
| I/σI | 10.53 (2.05) |
| Completeness % | 99.9 (99.9) |
| Redundancy | 5.7 (3.8) |
| **Refinement** |  |
| Resolution (Å) | 2.7 |
| No. unique reflections | 9,820 (454) |
| $R_{\text{work}}/R_{\text{free}}$ | 0.2452/0.3055 |
| No. atoms | 2,465 |
| Protein | 2,427 |
| Mg | 1 |
| Water | 37 |
| B-factors |  |
| Protein | 40.94 |
| Mg | 34.08 |
| R.m.s deviations |  |
| Bond lengths (Å) | 0.003 |
| Bond angles (°) | 0.8 |
| Ramachandran plot (%) |  |
| Favored | 90.76 |
| Allowed | 6.93 |
| Disallowed | 2.31 |

\* Highest-resolution shell is shown in parentheses.

**Figure 1.**
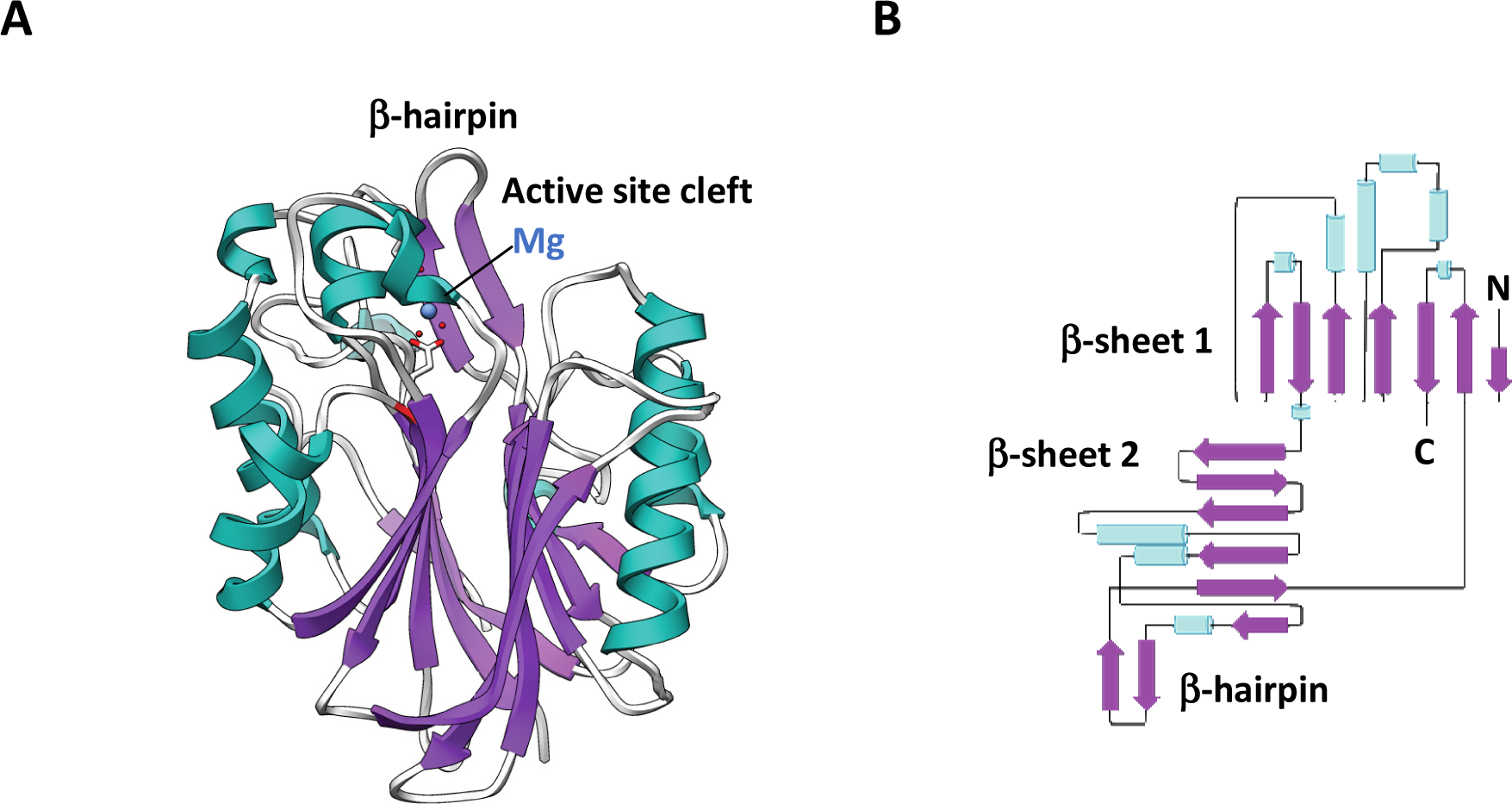
NOCT structure overview. **A)** Ribbon representation of human NOCT catalytic domain.**A** single magnesium ion shown is located in the catalytic center. **B)** Topologic connectivity diagram of NOCT with secondary structure elements shown.

**Figure 2.**
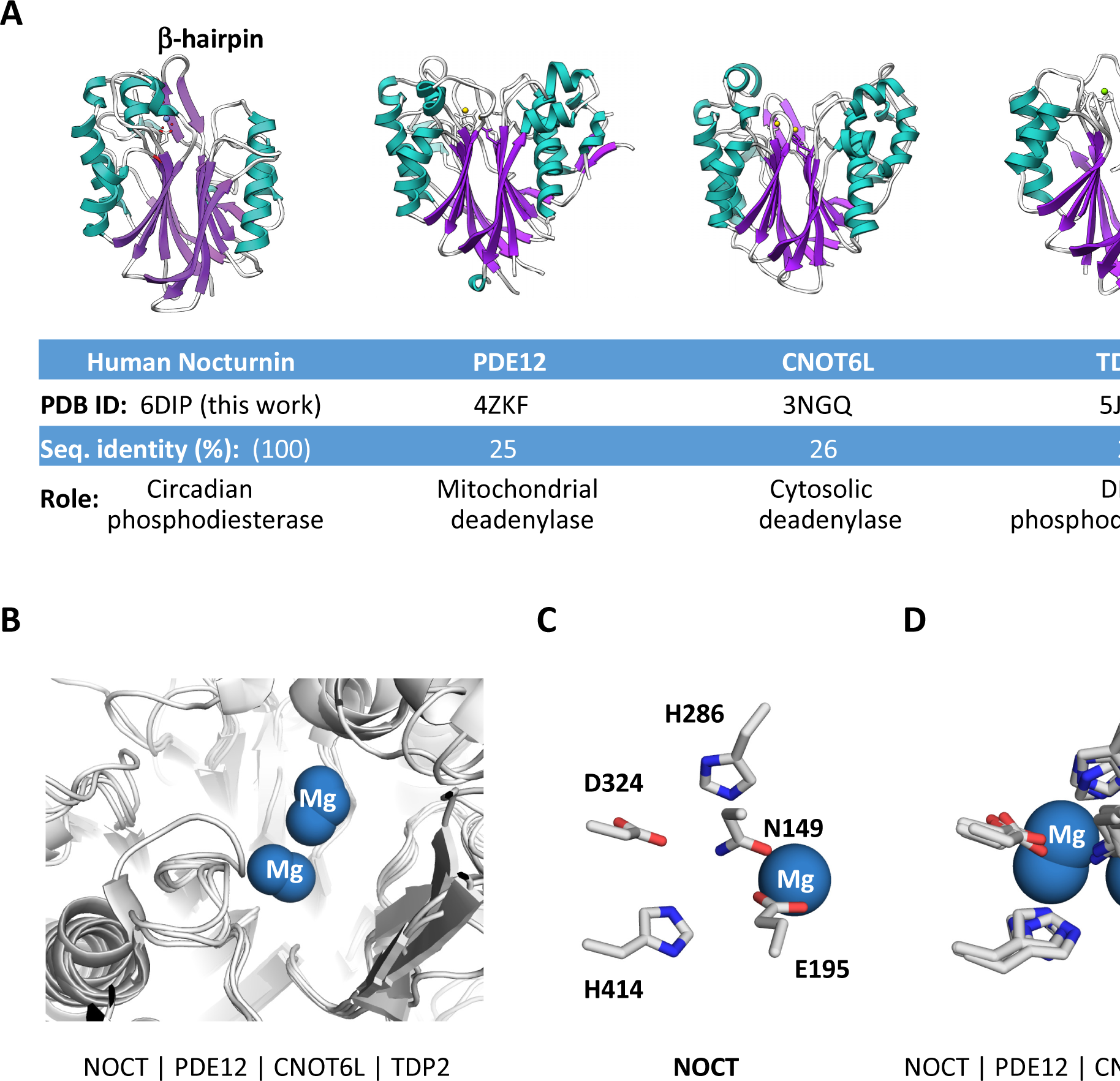
Structural relationships between NOCT and related phosphodiesterases. **A)** Structures, sequence identity and function summary for NOCT, PDE12, CNOT6L and TDP2. **B)** Active site superposition for NOCT, PDE12, CNOT6L and TDP2. **C)** Catalytic residues in the active site of human NOCT. **D)** Superposition of the catalytic residues in RNA and DNA hydrolases. All residues are numbered using human NOCT as a reference.

Further extending the analogy with the previously characterized magnesium-dependent phosphodiesterases, NOCT has a magnesium ion bound in the active site (Fig. 2B). All the key catalytic residues that coordinate magnesium and participate in RNA or DNA cleavage by α/β sandwich hydrolases are present in NOCT and occupy the same positions as in the previously described enzymes (Fig. 2C-D). Our structure therefore supports the model that NOCT mediates circadian function by acting as an enzyme hydrolyzing a phosphodiester bond.

### NOCT has a unique surface character and a surprisingly narrow active site

Although the secondary structure of NOCT is similar to those of PDE12, CNOT6L and TDP2, the surface properties of NOCT have a number of unique features. The global electrostatics of NOCT is distinct from that of the deadenylases PDE12 and CNOT6L due to the presence of a vast acidic area and a vast basic area near the active site in NOCT (Fig. 3A). Neither PDE12 nor CNOT6L have these areas and their electrostatics on the active site face arises largely from the acidic residues in the catalytic center. Although the electrostatic properties of NOCT and TDP2 are different, both proteins have patches of positive charge in similar locations. Surface charge properties of human NOCT are therefore more closely related to TDP2 than to PDE12/CNOT6L (Fig. 3A).

**Figure 3.**
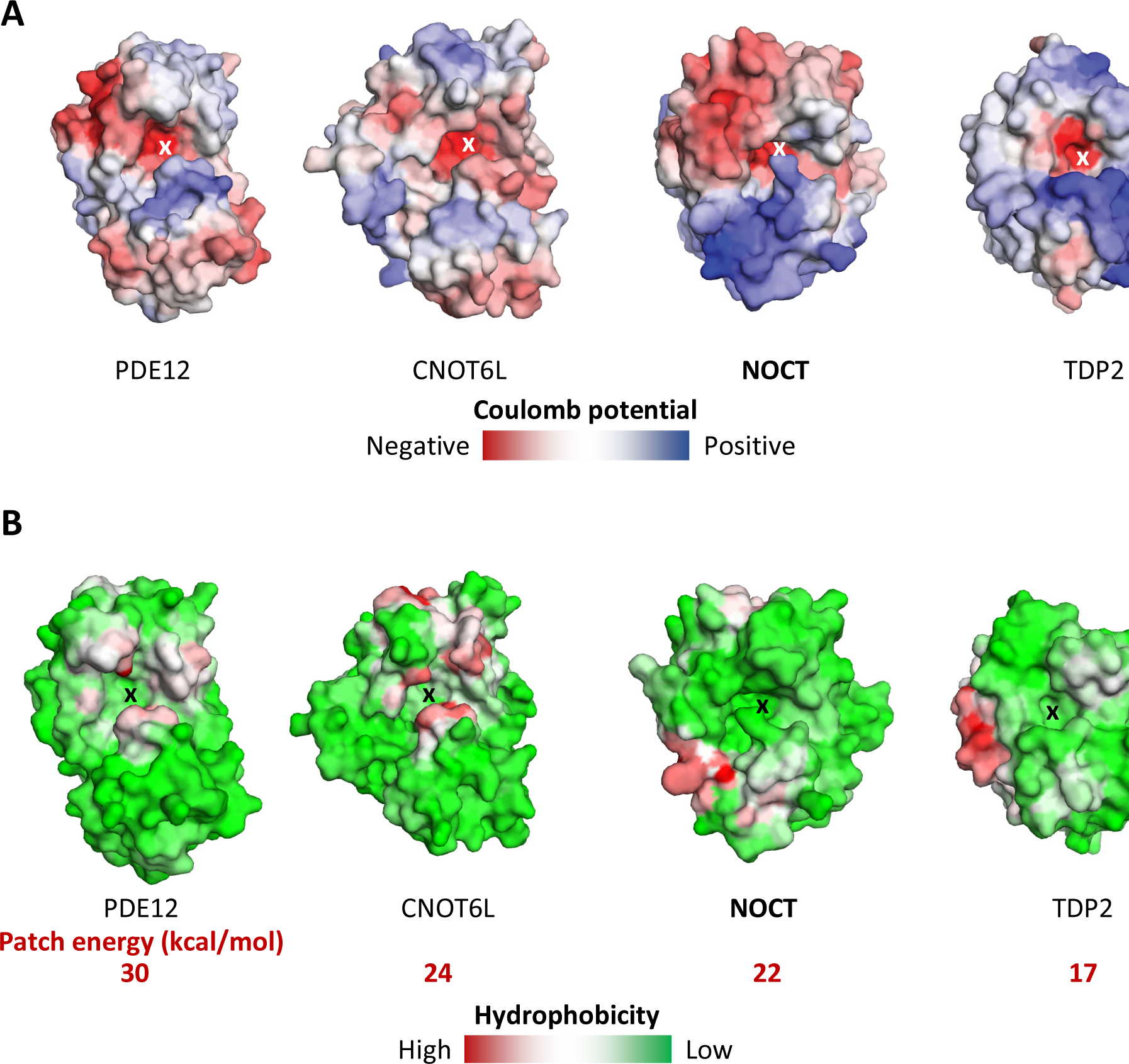
Surface analysis of NOCT and related phosphodiesterases. **A)** Vacuum electrostatic potential calculated using Coulomb’s law in SEQMOL-Kd (Methods). **B)** Surface area hydrophobic potential calculated using pre-atom desolvation analysis SEQMOL-Kd. The scale of red color (maximum hydrophobicity) in kcal/mol is shown for each protein.

Further differences between NOCT and the canonical deadenylases were revealed upon analysis of the hydrophobic potentials. The deadenylases PDE12/CNOT6L have similar hydrophobic potential configurations, with hydrophobic hot spots flanking the catalytic centers (Fig. 3B). These hot spots likely provide the interaction energy for recognition of nucleobases in poly-A RNA substrates. In contrast, NOCT lacks these hydrophobic hot spots altogether (Fig. 3B). The hydrophobic potential of NOCT resembles that of TDP2 more closely than those of PDE12 and CNOT6L: only NOCT and TDP2 are devoid of the hydrophobic patches around the catalytic core. Moreover, both NOCT and TDP2 evolved a hydrophobic patch on the surface located to the left of the active site (Fig. 3B), which could function as a docking site for binding partner proteins.

Analysis of active site accessibility reveals an additional difference between NOCT and the other α/β sandwich hydrolases. The active site pocket in NOCT is slightly longer than that in the other enzymes (Fig. 4A). Unexpectedly, the active site appears to be closed by a lid created by residue R290 (Fig. 4A-B). The narrow space created by the placement of R290 appears incompatible with binding of RNA, suggesting that R290 may have to move to permit substrate entry.

**Figure 4.**
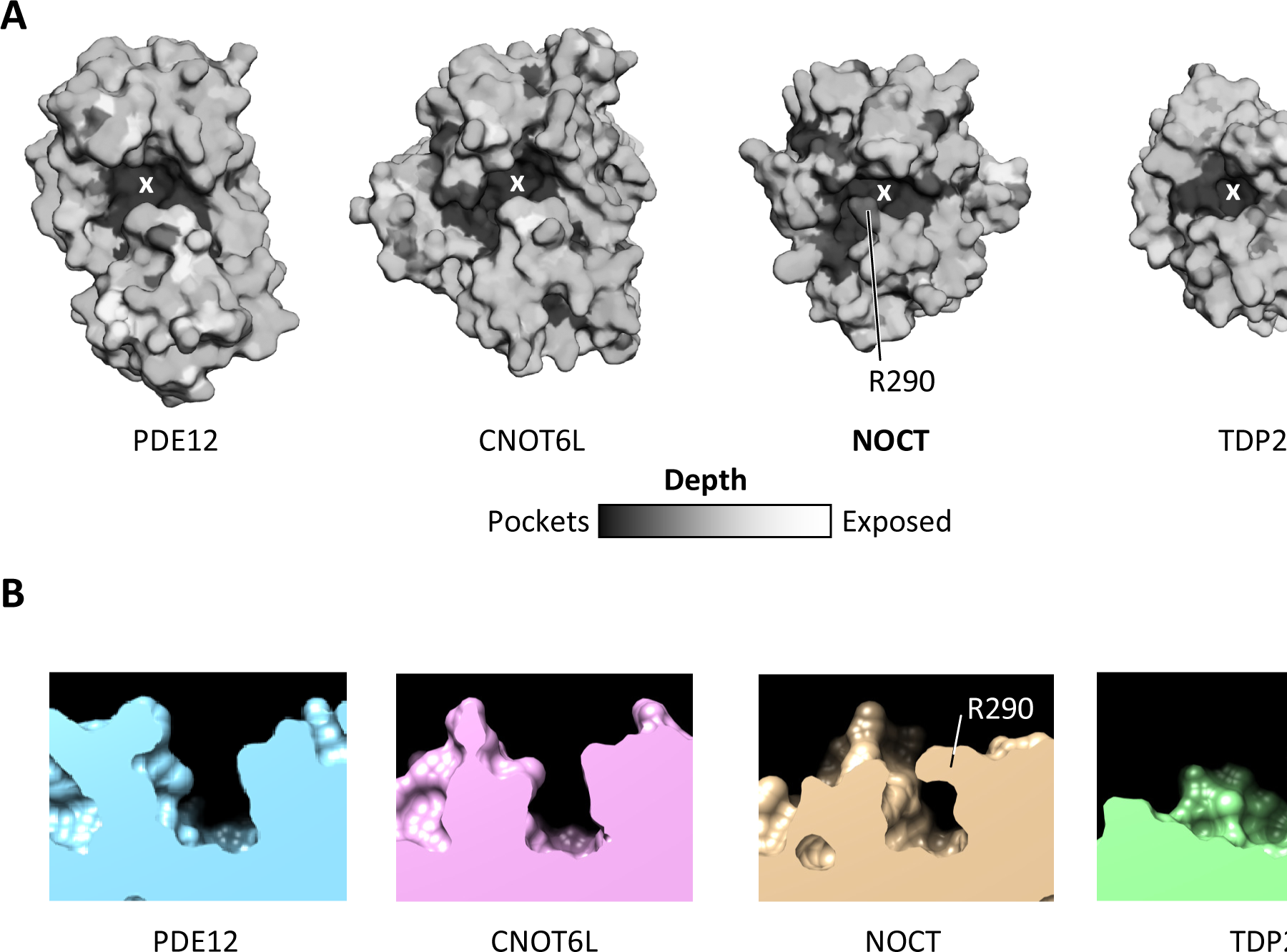
Active site properties of NOCT. **A)** Accessibility for a probe of 2.5 Å radius calculated in SEQMOL-Kd. Darker areas show deeper pockets. **B)** Side view (slice) of the active sites in NOCT, PDE12, CNOT6L and TDP2. All proteins were superimposed and positioned identically on the panels. TDP2 has the most open active site, whereas access to the NOCT active site is occluded by R290.

### Conservation analysis reveals unique structural elements near the active site present only in NOCT

Sequence conservation analysis remains one of the most reliable methods to attribute functional importance to protein residues. To obtain the conservation data, we identified 351 non-redundant NOCT sequences^10^ and carried out conservation analysis of NOCT crystal structure using SEMOL-Kd (Methods). This analysis revealed a strong conservation in the active site of NOCT, including conservation of the lid formed by the residue R290 (Fig. 5A). Therefore, conservation of the catalytic residues and phosphodiesterase activity is important for NOCT function. Moreover, the residue R290 is validated as a conserved part of the NOCT active site that has a yet unknown function.

**Figure 5.**
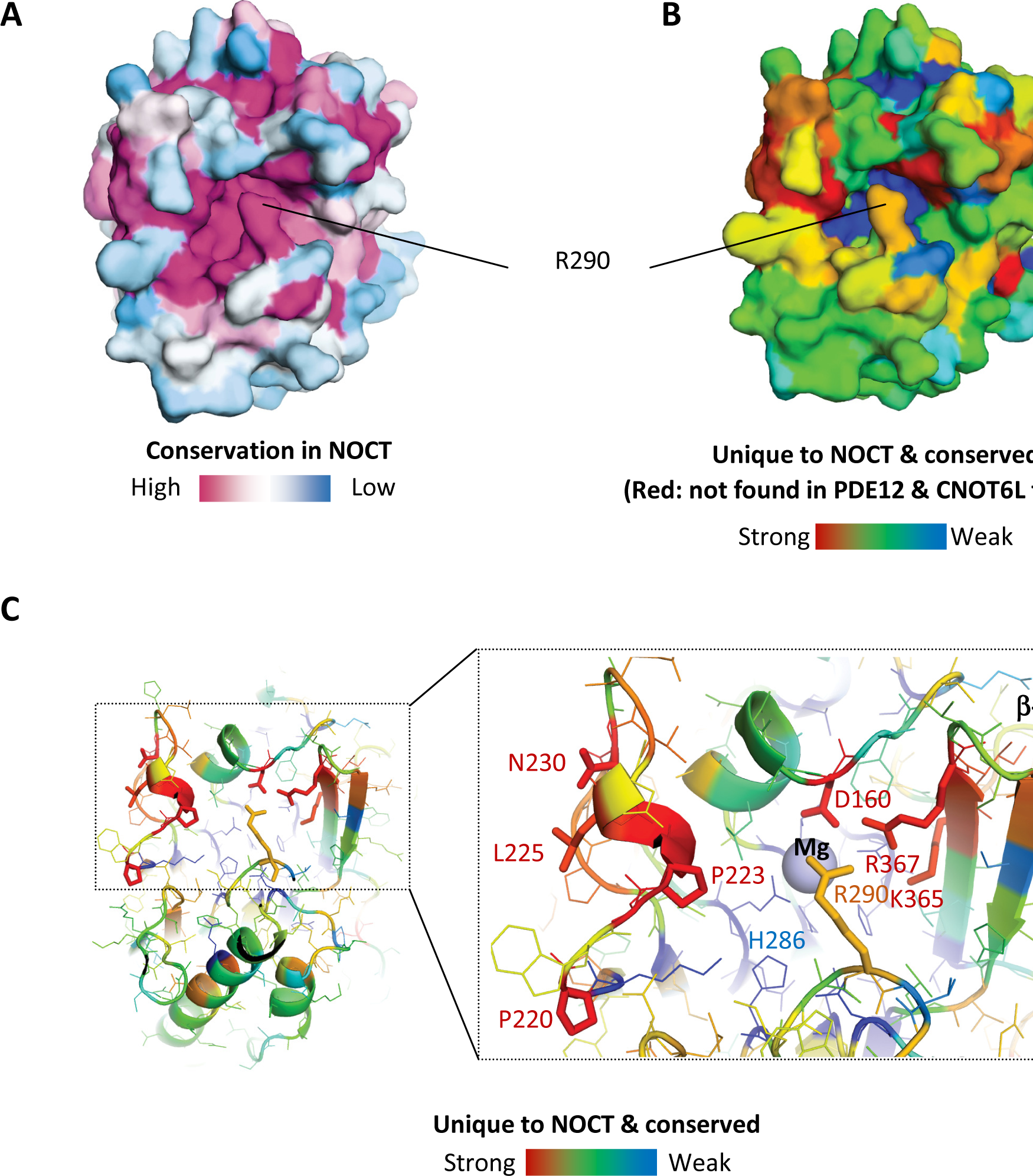
Conservation analysis for NOCT. **A)** NOCT conservation calculated from 351 non-redundant protein sequences using SEQMOL-Kd. Multiple sequence alignment was conducted with built-in Muscle 3.7^12^. **B-C)** Modified conservation analysis to visualize residues present only in NOCT family members, but absent in the families of PDE12 and CNOT6L deadenylases. Multiple NOCT-specific residues surround the active site.

Considering that NOCT has been described as a deadenylase^3,4^, we extended the conservation mapping analysis to allow straightforward comparisons of NOCT with PDE12 and CNOT6L. To this end we obtained 339 non-redundant sequences of PDE12 and 785 non-redundant sequences of CNOT6L. We calculated the net conservation scores for all residues in PDE12 and CNOT6L and arranged the scores in register with the NOCT amino acids. Next, for each NOCT residue we calculated the parameter S as follows:

S = ([NOCT_conservation] •[1-[PDE12&CNOT6L_conservation]])^0.5^

This modified conservation score (S) will attribute low values to residues conserved in NOCT, PDE12 and CNOT6L, such as catalytic amino acids. The low score will also be attributed to residues that are not conserved in NOCT. High score will be attributed only to NOCT residues that are (i) conserved in the NOCT family and (ii) are not conserved in the PDE12 and CNOT6L families. The resulting image reveals the structural elements that are both unique and invariable in NOCT (Fig. 5B, red). A close-up view of the S-colored PDB structure shows that in addition to the residue R290, NOCT harbors multiple conserved NOCT-specific residues around the active site. These residues include P220, P223, L225 and N230 in the coil to the left of the active site, D160 in the alpha-helix-loop motif above the active site, and K365 and R367 in the β-hairpin (Fig. 5C).

## Discussion

Our structure of human NOCT reveals the expected resemblance with the deadenylases PDE12 and CNOT6L, and a close resemblance with the DNA repair enzyme TDP2. The electrostatic properties and especially the hydrophobic properties of NOCT are nevertheless considerably divergent. We found that NOCT does not have the hydrophobic hot spots surrounding the catalytic pocket, which are apparently involved in poly-A nucleotide docking to PDE12 and CNOT6L. Presently we do not fully understand how NOCT recognizes RNA without these hydrophobic areas.

We found further that the active site of NOCT is occluded by the residue R290. Whereas side chains often can move and R290 could potentially change conformation upon RNA docking, conformational changes have energetic costs. The required free energy will inevitably weaken the affinity for RNA. However, conformational changes can provide the advantage because they can act as switches regulating the catalytic activity. The evolutionarily importance of R290 and the expectation that R290 has to move to allow RNA entry suggest that NOCT could be a regulated, rather than a constitutively active enzyme. Our efforts to co-crystallize NOCT with poly-A and poly-dA, including tests with NOCT catalytic mutants, produced only apo crystals. The interference with R290 could be one of the explanations for the difficulty of capturing NOCT with nucleic acids bound. Further understanding of NOCT must await the availability of a structure between NOCT and its cognate substrate.

## Materials and methods

### Cloning

The coding region of full-length human Nocturnin (1-431) was amplified by PCR from an in-house cDNA library using poly-IC transfected A549 cells and cloned into pGEX-6P vector (GE Healthcare Life Sciences). The Nocturnin 122-431 construct was made using site-directed mutagenesis that deleted DNA base pairs corresponding to residues 1-121. All constructs used in this study were verified by DNA sequencing.

### Protein Purification

pGEX-6P vector containing full-length human Nocturnin was transformed into *Escherichia coli* BL21 (DE3)-CodonPlus RIPL (Agilent Technologies) and grown to an OD_600_ of 0.4 in Luria-Bertani medium at 37 °C followed by induction with 0.2 mM isopropyl-β-D-thiogalactopyranoside (IPTG) and overnight expression at 22 °C. The cells were pelleted at 4,600 × *g* for 20 min, resuspended in lysis buffer [20 mM HEPES (pH 7.4), 1 M KCl, 2 mM MgCl_2_, 1 mM EDTA, 10% (vol/vol) glycerol, 5 mM DTT, and 1× Roche Complete protease inhibitors], and lysed on an EmulsiFlex C3 (Avestin). Crude lysates were clarified by centrifugation at 35,000 × *g* for 30 min, at 4 °C. Clarified lysates were affinity-purified using glutathione Sepharose (GE Healthcare Life Sciences) and the GST tag was removed with Prescission protease (GE Healthcare Life Sciences). Full-length Nocturnin was further purified by MonoQ, then MonoS ion-exchange chromatography, and lastly by Superdex 200 size-exclusion chromatography. Nocturnin 122-431 was further purified using a slight variation of performing MonoS first, then MonoQ ion-exchange chromatography, followed by Superdex 200 size-exclusion chromatography. All proteins were purified to more than 95% purity and concentrations were quantified by UV spectrophotometry.

### Nocturnin 122-431 Crystallization

Crystallization drops of Nocturnin 122-431 contained 17 mg/mL of protein and crystals were grown using the hanging drop vapor diffusion method by mixing the crystallization complex 1:1 with reservoir solution (0.1M MgCl_2_, 0.1M HEPES-Na pH 7.5, 30% PEG 400). Crystals were directly frozen in liquid nitrogen.

### X-Ray Data Collection and Structure Determination

X-ray diffraction data were collected using our Core facility Rigaku MicroMax-007 HF rotating anode generator supplied with Pilatus3 R 300K hybrid pixel array detector. Data were collected at a wavelength of 1.54 Å. Data were processed with the XDS package. Crystals contain one Nocturnin 122-431 molecule in the asymmetric unit and belong to the tetragonal P4_1_2_1_2 space group. The structure was solved by molecular replacement in PHASER using human CNOT6L (PDB ID code 3NGO) as the search model. The structure was modeled in COOT and refined by simulated annealing using PHENIX.

### Structure visualization and analysis

Structures were visualized with PyMol (DeLano Scientific build) and UCSF Chimera^11^. Surface properties (conservation, hydrophobic potential, electrostatics, pocket accessibility) were calculated and mapped using SEQMOL-Kd (BiochemLabSolutions, http://biochemlabsolutions.com/FASTAandPDB.html).

### Database entries

Structure factors and coordinates were deposited to PDB database (RCSB.org) under accession ID 6DIP.

## Acknowledgments

The authors would like to thank Dr. Phil Jeffrey (Princeton University) for the great help at the in house X-ray facility.

## References

1. Baggs, J.E. & Green, C.B. Functional analysis of nocturnin: a circadian clock-regulated gene identified by differential display. Methods Mol Biol 317, 243–54 (2006).

2. Gronke, S., Bickmeyer, I., Wunderlich, R., Jackle, H. & Kuhnlein, R.P. Curled encodes the Drosophila homolog of the vertebrate circadian deadenylase Nocturnin. Genetics 183, 219–32 (2009).

3. Baggs, J.E. & Green, C.B. Nocturnin, a deadenylase in Xenopus laevis retina: a mechanism for posttranscriptional control of circadian-related mRNA. Curr Biol 13, 189–98 (2003).

4. Stubblefield, J.J. et al. Temporal Control of Metabolic Amplitude by Nocturnin. Cell Rep 22, 1225–1235 (2018).

5. Pearce, S.F. et al. Maturation of selected human mitochondrial tRNAs requires deadenylation. Elife 6(2017).

6. Wang, H. et al. Crystal structure of the human CNOT6L nuclease domain reveals strict poly(A) substrate specificity. EMBO J 29, 2566–76 (2010).

7. Freudenthal, B.D., Beard, W.A., Cuneo, M.J., Dyrkheeva, N.S. & Wilson, S.H. Capturing snapshots of APE1 processing DNA damage. Nat Struct Mol Biol 22, 924–31 (2015).

8. Holm, L. & Rosenstrom, P. Dali server: conservation mapping in 3D. Nucleic Acids Res 38, W545–9 (2010).

9. Hornyak, P. et al. Mode of action of DNA-competitive small molecule inhibitors of tyrosyl DNA phosphodiesterase 2. Biochem J 473, 1869–79 (2016).

10. Gasteiger, E. et al. ExPASy: The proteomics server for in-depth protein knowledge and analysis. Nucleic Acids Res 31, 3784–8 (2003).

11. Pettersen, E.F. et al. UCSF Chimera--a visualization system for exploratory research and analysis. J Comput Chem 25, 1605–12 (2004).

12. Edgar, R.C. MUSCLE: multiple sequence alignment with high accuracy and high throughput. Nucleic Acids Res 32, 1792–7 (2004).

